# High Frequency Oscillations (>250Hz) Outnumber Interictal Spikes in Preclinical Studies of Alzheimer’s Disease

**DOI:** 10.1101/2023.10.30.564797

**Authors:** Christos Panagiotis Lisgaras, Helen E. Scharfman

## Abstract

Interictal spikes (IIS) and seizures are well-documented in Alzheimer’s disease (AD). IIS typically outnumber seizures, supporting their role as a prominent EEG biomarker in AD. In preclinical models, we showed that high frequency oscillations (HFOs>250Hz) also occur, but it is currently unknown how HFOs compare to IIS. Therefore, we asked whether the incidence of HFOs and IIS differed and if they are differentially affected by behavioral state.

We used three mouse lines that simulate aspects of AD: Tg2576, presenilin 2 knockout, and Ts65Dn mice. We recorded and quantified HFOs and IIS in the hippocampus during wakefulness, slow-wave sleep, and rapid eye movement sleep.

In all three mouse lines, HFOs were more frequent than IIS. High numbers of HFOs correlated with fewer IIS, suggesting for the first time possible competing dynamics among them in AD. Notably, HFOs occurred in more behavioral states than IIS.

In summary, HFOs were the most abundant EEG abnormality when compared to IIS, and occurred in all behavioral states, suggesting they are a better biomarker than IIS. These findings pertained to three mouse lines, which is important because they simulate different aspects of AD. We also show that HFOs may inhibit IIS.

**SHORT SUMMARY:** Interictal spikes (IIS) and seizures are common in Alzheimer’s disease (AD). IIS are more frequent than seizures and occur during earlier disease stages. In preclinical models, we showed that high frequency oscillations (HFOs>250Hz) occur, but a comparison between IIS and HFOs is lacking. Here we used 3 mouse lines with AD features and local field potential recordings to quantify IIS and HFOs. We found that HFOs outnumbered IIS and that their total numbers were inversely correlated with IIS. HFOs occurred during more behavioral states than IIS. Therefore, HFOs were the most abundant EEG abnormality, and this was generalizable across 3 types of preclinical AD.

## INTRODUCTION

Abnormal electrical activity strongly impacts outcomes in Alzheimer’s disease (AD). Indeed, AD patients with interictal spikes (IIS) show faster cognitive decline than those without IIS.^1^ Furthermore, the occurrence of seizures in patients with AD has been suggested to accelerate disease progression.^2,3^ Indeed, in rodents, seizures lead to amyloid-β (Aβ) release into the extracellular space, promoting deposition.^4^ However, seizures in AD can be rare^5-7^ and may accompany advanced disease stages whereas IIS occur earlier.^8,9^ Thus, a focus on interictal activity is warranted from a diagnostic and therapeutic standpoint.^10^

IIS are found in a subset of patients with AD (ranging from 22% to 53%)^1,8^ and patients with IIS that are treated with antiseizure medications show improved memory.^11^ Also, IIS are considered useful as a biomarker because they correlate with seizures in AD especially those that occur during rapid-eye movement (REM) sleep or wakefulness.^8^ In preclinical models, we showed that high frequency oscillations (HFOs) occur in 3 mouse lines that simulate AD features.^12^ Similar to IIS,^13-15^ HFOs occurred primarily during sleep, and they were also detectable early during disease progression.^12^ Nevertheless, a direct comparison between the rate of HFOs and IIS is lacking. This is an important gap in knowledge because new and improved EEG biomarkers are needed, and the best biomarker would be the one that is most common. Also, if the biomarker occurs in all behavioral states that would render it more efficient because a routine EEG would capture it.

Here we focused on determining how HFOs compare with IIS. We addressed the following questions: Do HFOs occur more frequently than IIS? Do HFOs occur in all behavioral states and outnumber IIS in all states? We also asked whether the total number of IIS and HFOs correlated because such data would uncover potential synergistic or competing dynamics between them.

## MATERIALS AND METHODS

### I. Animals

We used 3 mouse lines that we previously reported exhibit HFOs^12^ and IIS.^13,16,17^ In brief, Tg2576 mice (bred in-house^13,18^) overexpress a mutated form of the human amyloid precursor protein simulating a Swedish family with AD using the hamster prion promoter (APPSwe;^13,18^). They were used as a model of progressive Aβ overexpression.^13,18-20^ A total of 8 Tg2576 transgenic mice (4 males, 4 females) and 5 Tg2576 control mice (3 males, 2 females) were used. At the time of recording, Tg2576 transgenic mice were 5.1±1.9 months-old and controls were 5.3±3.1 months-old (Mann-Whitney *U*-test, U=25, p>0.99). Presenilin (PS2) knockout (PS2KO) mice were used to simulate reduced function of PS2 similar to mutations in *PS2* gene seen in a form of familial early onset AD.^21^ We used 4 PS2KO (2 males, 2 females) and 4 controls (2 males, 2 females). PS2KO mice were 10.7±1.9 months-old and controls were 10.7±1.9 months-old (unpaired t-test, t_*crit*_=0.009, df=6, p=0.99). The Ts65Dn mouse model of Down syndrome (stock #005252, The Jackson Laboratory) was used since the majority of individuals with Down syndrome develop AD.^22-24^ These mice have a triplicated portion of the mouse chromosome 16 which is the ortholog of the human chromosome triplicated in individuals with Down syndrome.^25^ We used 4 transgenic (2 males, 2 females) and 5 control (3 males, 2 females) mice. Ts65Dn transgenic mice were 13.2±2.9 months-old and controls were 15.3±4.6 months-old (Mann-Whitney *U*-test, U=24.5, p=0.76). There were no significant differences in age between the controls of different mouse lines (Kruskal-Wallis test, H(3)=3.89, p=0.14) nor the transgenic or knockout mice (Kruskal-Wallis test, H(3)=5.96, p=0.06). Also, we found no sex differences in IIS rate during REM (Mann-Whitney *U*-test, U=31.5, p=0.45); please note that IIS were not detected during wakefulness or slow-wave sleep (SWS) thus no sex differences could be studied during these behavioral states. We found no sex differences in HFO rate during wakefulness (unpaired t-test, t_*crit*_=1.03, df=14, p=0.32), SWS (unpaired t-test, t_*crit*_=0.81, df=14, p=0.43), REM sleep (unpaired t-test, t_*crit*_=0.79, df=14, p=0.44) or total number of HFOs in all behavioral states (unpaired t-test, t_*crit*_=0.85, df=14, p=0.85). In light of no apparent sex differences, in all analyses, we pooled together data from male and female mice.

Tg2576 mice were bred on a C57BL6/SJL background (stock #100012, The Jackson Laboratory), PS2KO on a C57BL/6J background (stock #000664, The Jackson Laboratory) and Ts65Dn mice on a hybrid background (stock #003647, The Jackson Laboratory). For breeding, wild type female mice were bred with Tg2576 transgenic or PS2KO male mice. Breeding for Ts65Dn mice used a transgenic Ts65Dn female mice crossed to a wild type male mice. The breeders were fed Purina 5008 and after weaning, mice were fed Purina 5001 (all food from W.F. Fisher) with water ad libitum. They were housed with the same sex (<4 mice/cage) in cages with corn cob bedding and there was a 12hr light:dark cycle (7:00 a.m. lights on, 7:00 p.m. lights off). Genotyping was performed by the Mouse Genotyping Core Laboratory at New York University Langone Medical Center. All experimental procedures were performed in accordance with the NIH guidelines and approved by the Institutional Animal Care and Use Committee at the Nathan Kline Institute.

### II. Electrode implantation and recordings

Mice were anesthetized, placed in a stereotaxic apparatus, and buprenorphine (0.05mg/kg, s.c.) was injected to reduce discomfort. Two burr holes were drilled over the cerebellum and reference and ground screw electrodes were stabilized there using dental cement (Lang Dental). Next, one 16-channel silicon probe (#A1x16, Neuronexus or #PLX-QP-3-16E-v2, Plexon) or a single wire (90μm diameter stainless steel, California Fine Wire) was implanted in the left dorsal DG (- 1.9mm A-P, -1.2mm M-L, -1.9mm D-V). Local field potential signals were recorded at 2kHz using a bandpass filter (0.1-500Hz) with either the Digital Lynx SX (Neuralynx), an RHD interface board (Intan Techologies), or Sirenia with simultaneous video (#ac2000, Basler). Previous studies showed that the different hardware all captured IIS and HFOs to the same extent, so data were pooled.^12,16^ Mice were continuously recorded for 3 consecutive days at a minimum. IIS and HFOs were quantified from the 2^nd^ 24hr-long period of the recording.

### III. IIS and HFO detection

IIS and HFOs were detected using the same criteria as described in our previous studies.^12,13,16,26,27^ We quantified IIS or HFOs that occurred during periods of wakefulness, SWS, or REM; these behavioral states were defined as before.^12,16^ IIS were included if their amplitude exceeded the mean of the baseline noise by more than 5 standard deviations. This criterion was chosen because it excluded movement artifacts or other types of noise. We filtered the EEG in the 250-500Hz frequency band to detect HFOs, and used an automated approach in the RippleLab^28^ and the same criteria as in our previous HFO studies.^12,27^

### IV. Statistics

Data are presented as a mean ± SEM and statistical significance was set at p<0.05; non-significant comparisons are designated by “ns.” All statistical analyses were performed using Prism (Graphpad). The Shapiro-Wilk test was used to test if the data fit a normal distribution. Brown-Forsythe was used to test homogeneity of variance.

Comparisons of means for parametric data of two groups were conducted using paired or unpaired two-tailed t-tests; non-parametric data used Wilcoxon signed rank, or Mann-Whitney tests. For comparisons of more than 2 groups, ANOVA was used followed by *post-hoc* tests and corrections for multiple comparisons. Repeated measures ANOVA used a mixed-effects model because of missing data points, followed by Holm-Šídák’s *post-hoc* tests.

For non-parametric data with more than 2 groups, Kruskal-Wallis was used, and Dunn’s *post-hoc* test with correction for multiple comparisons. For correlation analyses, we used linear regression. The Pearson correlation coefficient (*r*) was used for parametric data and Spearman *r* for non-parametric data. A Chi-square test was used to compare the percentage of animals that showed HFOs and/or IIS during the different behavioral states.

## RESULTS

### I. HFOs are more frequent than IIS

We found that all transgenic and knockout mice showed IIS and HFOs (n=8 Tg2576, n=4 PS2KO, n=4 Ts65Dn). None of the 14 age-matched controls showed either IIS or HFOs (n=5 Tg2576, n=4 PS2KO, n=5 Ts65Dn). We next compared the total number of detected HFOs and IIS separately for each mouse line (Fig 1A). A mixed effects model with the total number of events (HFOs or IIS) and mouse line (Tg2576, PS2KO, or Ts65Dn) as factors showed a significant effect (Mixed effect model, F(1.54, 5.87) = 49.92, p=0.0003; Fig 1A). *Post-hoc* comparisons showed that the number of HFOs was significantly greater compared to IIS in every mouse line we used (Tg2576, p <0.0001; PS2KO, p=0.009; Ts65Dn, p=0.01; Fig 1A). Also, the dominance of HFOs over IIS (HFO/IIS ratio) depended on the mouse line (Kruskal-Wallis test, H(3) = 9.54, p=0.001; Fig 1B) and that is important because increased ratios of HFOs>250Hz are linked to pathophysiological alterations seen in epilepsy.^29^ Indeed, *post-hoc* comparisons showed a greater HFO/IIS ratio in Ts65Dn mice vs. Tg2576 (p=0.01; Fig 1B) mice but not PS2KO (p>0.99; Fig 1B) or between Tg2576 and PS2KO mice (p=0.05; Fig 1B).

**Figure 1:**
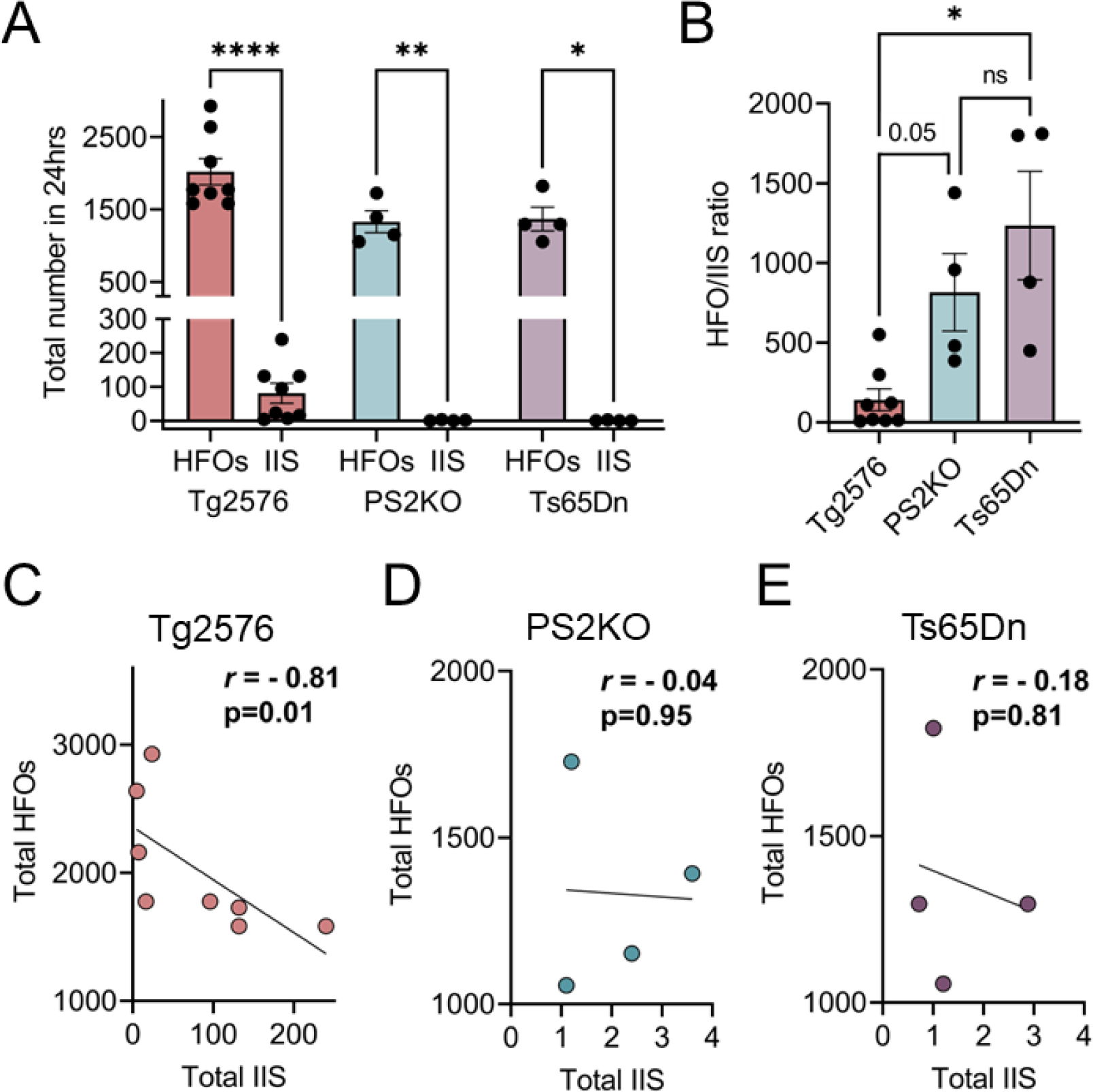
HFOs are more frequent than IIS and their rates are inversely correlated. A. The total number of HFOs or IIS is shown during 24hr in each mouse line. There was a significant effect between the total number of events and mouse line (Mixed effects model, F (1.54, 5.87) = 49.92, p=0.0003). *Post-hoc* comparisons showed that the total number of HFOs was significantly greater compared to IIS in every mouse line we used (Tg2576, p <0.0001; PS2KO, p=0.009; Ts65Dn, p=0.01). B. HFO to IIS ratio was significantly different between mouse lines (Kruskal-Wallis test, H(3) = 9.54, p=0.001). *Post-hoc* comparisons showed a greater HFO/IIS ratio in Ts65Dn mice vs. Tg2576 (p=0.01) mice but not PS2KO (p>0.99) or between Tg2576 and PS2KO (p=0.05). C. The number of HFOs was inversely correlated to the number of IIS in Tg2576 mice (Spearman *r* = -0.81, p=0.01). D. Lack of linear correlation between HFOs and IIS in PS2KO mice (Pearson *r* = -0.04, p=0.95). E. Same as in D but for Ts65Dn mice. We found a lack of linear correlation between the total number of HFOs and IIS (Pearson *r* = -0.18, p=0.81).

### II. HFOs are inversely correlated with IIS

Next, we asked whether the greater number of HFOs would correlate with more frequent IIS. Instead, we found that the number of HFOs was inversely correlated with the number of IIS in Tg2576 mice (Spearman *r* = -0.81, p=0.01; Fig 1C). PS2KO mice did not show a linear correlation between the number of HFOs and IIS (Pearson *r* = -0.04, p=0.95; Fig 1D) similar to Ts65Dn mice (Pearson *r* = -0.18, p=0.81; Fig 1E). However, it should be noted that this lack of linear correlation may be due to few IIS in these two mouse lines.^16^

### III. HFOs are detectable during more behavioral states than IIS

We next asked whether HFOs and IIS occurred during the same behavioral states and if their rates differed according to behavioral state. First, we found that the percent of animals that showed HFOs or IIS was different across behavioral states (Chi-square test, p<0.0001; Fig 2A). Indeed, HFOs occurred during wakefulness, SWS, and REM in all mice we tested (16/16 mice). In contrast, IIS were only found to occur during REM (16/16 mice).

**Figure 2:**
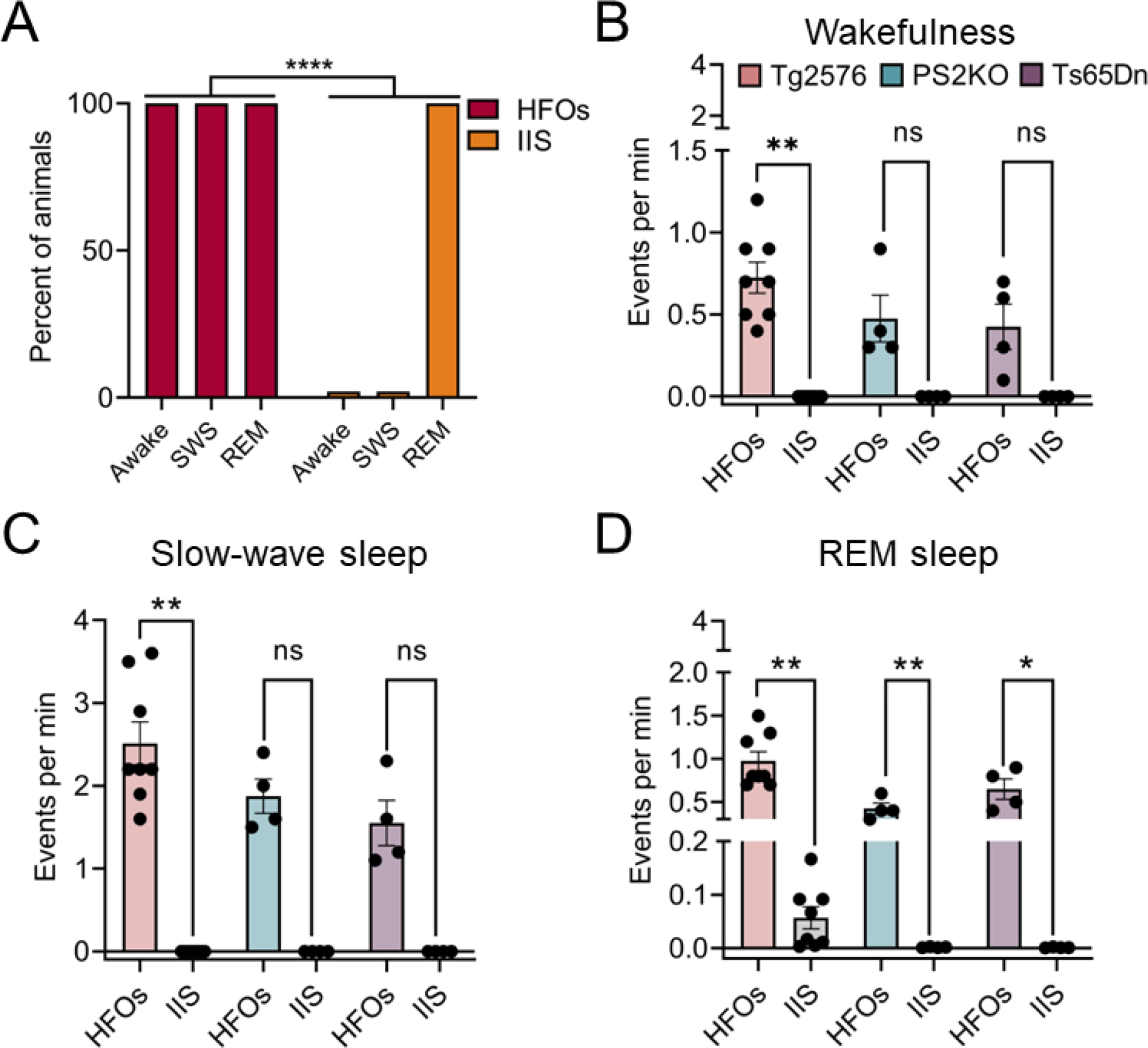
HFOs accompany more behavioral states than IIS and are more frequent regardless of behavioral state. A. The percent of animals that showed HFOs or IIS differed across behavioral states (Chi-square test, p <0.0001). All 16 mice (n=8 Tg2576, n=4 PS2KO, n=4 Ts65Dn) showed HFOs during wakefulness, SWS, and REM sleep. In contrast, IIS were only found to occur during REM (16/16 mice), but not during wakefulness (0/16 mice) or SWS (0/16 mice). B. HFO and IIS rates during wakefulness. The rate of HFOs was significantly higher than IIS in Tg2576 mice (Wilcoxon signed rank test, p=0.007) but not PS2KO (Wilcoxon signed rank test, p=0.12) or Ts65Dn mice (Wilcoxon signed rank test, p=0.12). C. HFO and IIS rates during SWS. Similar to wakefulness, we found that Tg2576 mice showed a significantly higher rate of HFOs vs. IIS (Wilcoxon signed rank test, p=0.007); PS2KO and Ts65Dn mice did not show a significant difference between HFO and IIS rate (PS2KO, Wilcoxon signed rank test, p=0.12; Ts65Dn, Wilcoxon signed rank test, p=0.12). D. HFOs and IIS during REM sleep. HFOs were more frequent than IIS in all mouse lines we used (Tg2576, Wilcoxon signed rank test, p=0.007; PS2KO, paired t-test, t_*crit*_=6.77, df=3, p=0.006; Ts65Dn, paired t-test, t_*crit*_=5.45, df=3, p=0.01).

We next compared the rate of IIS and HFOs separately for each behavioral state to better understand how their relationship may differ. During wakefulness, the rate of HFOs was significantly higher than IIS in Tg2576 mice (Wilcoxon signed rank test, p=0.007; Fig 2B) but not PS2KO (Wilcoxon signed rank test, p=0.12; Fig 2B) or Ts65Dn mice (Wilcoxon signed rank test, p=0.12; Fig 2B). During SWS, we found a similar difference in Tg2576 mice (Wilcoxon signed rank test, p=0.007; Fig 2C) but not in the other 2 mouse lines (PS2KO, Wilcoxon signed rank test, p=0.12; Ts65Dn, Wilcoxon signed rank test, p=0.12; Fig 2C). During REM sleep, HFOs were more frequent than IIS in all mouse lines we used (Tg2576, Wilcoxon signed rank test, p=0.007; PS2KO, paired t-test, t_*crit*_=6.77, df=3, p=0.006; Ts65Dn, paired t-test, t_*crit*_=5.45, df=3, p=0.01; Fig 2D).

## DISCUSSION

### Summary of main findings

The goal of this study was to determine whether HFOs or IIS are more frequent EEG disturbances in mouse lines that simulate AD. In all 3 mouse lines, we found that HFOs outnumbered IIS. In Tg2576 mice, HFOs appeared to inversely correlate with the number of IIS. Notably, more behavioral states showed HFOs vs. IIS. Last, we found that HFOs in Tg2576 occurred more frequently than IIS regardless of behavioral state. These results support the idea that HFOs are a prominent EEG abnormality in mice that simulate AD and suggest they could be a better biomarker than IIS.

### HFOs as a more abundant EEG abnormality than IIS

We found that HFOs occurred more frequently than IIS. In humans with AD, IIS occur in a fraction of patients and their rates may vary.^1^ Notably, we previously found that IIS rate in 3 mouse lines^16^ closely resembled those reported in humans with AD.^1^ If HFO rates in mice and humans are similar, then HFOs could be more abundant than IIS in humans and thus more likely to be captured. In epilepsy where HFOs also occur, humans and epileptic animals appear to share similar characteristics of HFOs, supporting their translational value.^30,31^ Nevertheless, future studies comparing HFOs and IIS in humans with AD are warranted to better understand whether they are more frequent than IIS.

### HFOs and IIS dynamics

We found that mice with more HFOs showed fewer IIS. In epileptic mice, it has been reported that an increased occurrence of HFOs correlated to impaired memory^32^ and that HFO disruption improves memory.^26^ Thus, it is likely that the increased number of HFOs we found may affect memory performance at least in PS2KO and Ts65Dn mice that IIS appear to be infrequent. However, it is important to also consider that IIS alone have detrimental effects on memory.^1,33,34^ Nevertheless, remains to be determined whether HFOs or IIS are more detrimental to memory in AD.

It is also noteworthy to consider that IIS occurring after an HFO in humans with epilepsy may show reduced neural firing.^35^ This reduced neural firing may make IIS less likely to generate because IIS represent a large synchronous discharge.^36^ If that is true in our mice as well, then frequent HFOs may have inhibited IIS.

### Potential limitations

PS2KO and Ts65Dn mice showed few IIS, and this might have led to an underestimation of correlations between HFOs and IIS. However, the rate of IIS we report is similar to human AD^1^ and that is notable from a translational perspective. Also, possible effects of age remain to be determined. However, the age ranges used in this study encompass a wide range and no clear differences obvious. This is based on our prior study of HFOs in these mouse lines which did not show an effect of age^12^ and our prior study of IIS showing that IIS occur at an extremely broad range of ages with again no apparent effect of age.^16^ Notably, at the ages included in this study, IIS rates in Tg2576 mice can show variability.^13,17^

## CONCLUSIONS

The striking predominance of HFOs over IIS is important because it makes HFOs a more promising biomarker. This is especially relevant for short EEG recordings conducted in an outpatient setting. In addition, because HFOs occur in all behavioral states they are better than IIS for an outpatient clinic. Although HFOs will require wide-band recording, these types of recordings are increasingly feasible. Because HFOs begin very early in the mouse lines and do so in three diverse lines, they hold great promise in identifying individuals at risk for AD at a very young age of life.

## Abbreviations

AD: Alzheimer’s disease
HFOs: High frequency oscillations
IIS: Interictal spike
NREM: Non-rapid eye movement sleep
PS2KO: Presenilin 2 knockout
REM: Rapid eye movement sleep
SWS: Slow-wave sleep

## ACKNOWLEDGEMENTS

This project was supported by NIH grants R01 AG-055328 and R01 NS-106983, and the New York State Office of Mental Health.

## CONFLICTS OF INTEREST/ETHICAL PUBLICATION STATEMENT

None of the authors has any conflict of interest to disclose. We confirm that we have read the Journal’s position on issues involved in ethical publication and affirm that this report is consistent with those guidelines.

## DATA AVAILABILITY STATEMENT

Data will be available from the corresponding author upon reasonable request.

## REFERENCES

1. Vossel KA, Ranasinghe KG, Beagle AJ, Mizuiri D, Honma SM, Dowling AF, Darwish SM, Van Berlo V, Barnes DE, Mantle M, Karydas AM, Coppola G, Roberson ED, Miller BL, Garcia PA, Kirsch HE, Mucke L, Nagarajan SS. Incidence and impact of subclinical epileptiform activity in Alzheimer’s disease. Ann Neurol. 2016;80(6):858–70.

2. Volicer L, Smith S, Volicer BJ. Effect of seizures on progression of dementia of the Alzheimer type. Dementia. 1995 Sep-Oct;6(5):258–63.

3. Förstl H, Burns A, Levy R, Cairns N, Luthert P, Lantos P. Neurologic signs in Alzheimer’s disease. Results of a prospective clinical and neuropathologic study. Arch Neurol. 1992 Oct;49(10):1038–42.

4. Cirrito JR, Yamada KA, Finn MB, Sloviter RS, Bales KR, May PC, Schoepp DD, Paul SM, Mennerick S, Holtzman DM. Synaptic activity regulates interstitial fluid amyloid-β levels in vivo. Neuron. 2005 Dec 22;48(6):913–22.

5. Hauser WA, Morris ML, Heston LL, Anderson VE. Seizures and myoclonus in patients with Alzheimer’s disease. Neurology. 1986 Sep;36(9):1226–30.

6. Scarmeas N, Honig LS, Choi H, Cantero J, Brandt J, Blacker D, Albert M, Amatniek JC, Marder K, Bell K, Hauser WA, Stern Y. Seizures in Alzheimer disease: who, when, and how common? Arch Neurol. 2009 Aug;66(8):992–7.

7. Friedman D, Honig LS, Scarmeas N. Seizures and epilepsy in Alzheimer’s disease. CNS Neurosci Ther. 2012 Apr;18(4):285–94.

8. Lam AD, Sarkis RA, Pellerin KR, Jing J, Dworetzky BA, Hoch DB, Jacobs CS, Lee JW, Weisholtz DS, Zepeda R, Westover MB, Cole AJ, Cash SS. Association of epileptiform abnormalities and seizures in Alzheimer disease. Neurology. 2020 Oct 20;95(16):e2259–e70.

9. Lam AD, Deck G, Goldman A, Eskandar EN, Noebels J, Cole AJ. Silent hippocampal seizures and spikes identified by foramen ovale electrodes in Alzheimer’s disease. Nat Med. 2017 Jun;23(6):678–80.

10. Sperling RA, Jack CR, Jr., Aisen PS. Testing the right target and right drug at the right stage. Sci Transl Med. 2011 Nov 30;3(111):111cm33.

11. Vossel K, Ranasinghe KG, Beagle AJ, La A, Ah Pook K, Castro M, Mizuiri D, Honma SM, Venkateswaran N, Koestler M, Zhang W, Mucke L, Howell MJ, Possin KL, Kramer JH, Boxer AL, Miller BL, Nagarajan SS, Kirsch HE. Effect of levetiracetam on cognition in patients with Alzheimer disease with and without epileptiform activity: a randomized clinical trial. JAMA Neurol. 2021 Nov 1;78(11):1345–54.

12. Lisgaras CP, Scharfman HE. High frequency oscillations (250-500Hz) in animal models of Alzheimer’s disease and two animal models of epilepsy. Epilepsia. 2022 Nov 8.

13. Kam K, Duffy AM, Moretto J, LaFrancois JJ, Scharfman HE. Interictal spikes during sleep are an early defect in the Tg2576 mouse model of β-amyloid neuropathology. Sci Rep. 2016 Jan 28;6:20119.

14. Brown R, Lam AD, Gonzalez-Sulser A, Ying A, Jones M, Chou RC, Tzioras M, Jordan CY, Jedrasiak-Cape I, Hemonnot AL, Abou Jaoude M, Cole AJ, Cash SS, Saito T, Saido T, Ribchester RR, Hashemi K, Oren I. Circadian and brain state modulation of network hyperexcitability in Alzheimer’s disease. eNeuro. 2018 Mar-Apr;5(2).

15. Gureviciene I, Ishchenko I, Ziyatdinova S, Jin N, Lipponen A, Gurevicius K, Tanila H. Characterization of epileptic spiking associated with brain amyloidosis in APP/PS1 mice. Front Neurol. 2019 2019-November-12;10.

16. Lisgaras CP, Scharfman HE. Interictal spikes in Alzheimer’s disease: Preclinical evidence for dominance of the dentate gyrus and cholinergic control by the medial septum. Neurobiol Dis. 2023 Sep 14;187:106294.

17. Chartampila E, Elayouby KS, Leary P, LaFrancois JJ, Alcantara-Gonzalez D, Jain S, Gerencer K, Botterill JJ, Ginsberg SD, Scharfman HE. Choline supplementation in early life improves and low levels of choline can impair outcomes in a mouse model of Alzheimer’s disease. bioRxiv. 2023 May 24.

18. Hsiao K, Chapman P, Nilsen S, Eckman C, Harigaya Y, Younkin S, Yang F, Cole G. Correlative memory deficits, Aβ elevation, and amyloid plaques in transgenic mice. Science. 1996 Oct 4;274(5284):99–102.

19. Ishii M, Wang G, Racchumi G, Dyke JP, Iadecola C. Transgenic mice overexpressing amyloid precursor protein exhibit early metabolic deficits and a pathologically low leptin state associated with hypothalamic dysfunction in arcuate neuropeptide Y neurons. J Neurosci. 2014 Jul 2;34(27):9096–106.

20. Kawarabayashi T, Younkin LH, Saido TC, Shoji M, Ashe KH, Younkin SG. Age-dependent changes in brain, CSF, and plasma amyloid β protein in the Tg2576 transgenic mouse model of Alzheimer’s disease. J Neurosci. 2001;21(2):372–81.

21. Lehmann L, Lo A, Knox KM, Barker-Haliski M. Alzheimer’s disease and epilepsy: a perspective on the opportunities for overlapping therapeutic innovation. Neurochem Res. 2021 Aug;46(8):1895–912.

22. Wisniewski KE, Wisniewski HM, Wen GY. Occurrence of neuropathological changes and dementia of Alzheimer’s disease in Down’s syndrome. Ann Neurol. 1985 Mar;17(3):278–82.

23. Salehi A, Ashford JW, Mufson EJ. The link between Alzheimer’s disease and Down syndrome. A historical perspective. Curr Alzheimer Res. 2016;13(1):2–6.

24. Head E, Powell D, Gold BT, Schmitt FA. Alzheimer’s disease in Down syndrome. Eur J Neurodegener Dis. 2012 Dec;1(3):353–64.

25. Reinholdt LG, Ding Y, Gilbert GJ, Czechanski A, Solzak JP, Roper RJ, Johnson MT, Donahue LR, Lutz C, Davisson MT. Molecular characterization of the translocation breakpoints in the Down syndrome mouse model Ts65Dn. Mamm Genome. 2011 Dec;22(11-12):685–91.

26. Lisgaras CP, Oliva A, McKenzie S, LaFrancois J, Siegelbaum SA, Scharman HE. Hippocampal area CA2 controls seizure dynamics, interictal EEG abnormalities and social comorbidity in mouse models of temporal lobe epilepsy. bioRxiv. 2023 Jan 18.

27. Lisgaras CP, Scharfman HE. Robust chronic convulsive seizures, high frequency oscillations, and human seizure onset patterns in an intrahippocampal kainic acid model in mice. Neurobiol Dis. 2022 Jan 25:105637.

28. Navarrete M, Alvarado-Rojas C, Le Van Quyen M, Valderrama M. RIPPLELAB: A comprehensive application for the detection, analysis and classification of high frequency oscillations in electroencephalographic signals. PLoS One. 2016;11(6):e0158276.

29. Staba RJ, Frighetto L, Behnke EJ, Mathern GW, Fields T, Bragin A, Ogren J, Fried I, Wilson CL, Engel Jr J. Increased fast ripple to ripple ratios correlate with reduced hippocampal volumes and neuron loss in temporal lobe epilepsy patients. Epilepsia. 2007;48(11):2130–8.

30. Bragin A, Engel J, Jr., Wilson CL, Fried I, Buzsaki G. High-frequency oscillations in human brain. Hippocampus. 1999;9(2):137–42.

31. Bragin A, Engel J, Jr., Wilson CL, Fried I, Mathern GW. Hippocampal and entorhinal cortex high-frequency oscillations (100-500 Hz) in human epileptic brain and in kainic acid-treated rats with chronic seizures. Epilepsia. 1999 Feb;40(2):127–37.

32. Valero M, Averkin RG, Fernandez-Lamo I, Aguilar J, Lopez-Pigozzi D, Brotons-Mas JR, Cid E, Tamas G, Menendez de la Prida L. Mechanisms for selective single-cell reactivation during offline sharp-wave ripples and their distortion by fast ripples. Neuron. 2017 Jun 21;94(6):1234-47.e7.

33. Kleen JK, Scott RC, Holmes GL, Lenck-Santini PP. Hippocampal interictal spikes disrupt cognition in rats. Ann Neurol. 2010 Feb;67(2):250–7.

34. Soula M, Maslarova A, Harvey RE, Valero M, Brandner S, Hamer H, Fernández-Ruiz A, Buzsáki G. Interictal epileptiform discharges affect memory in an Alzheimer’s disease mouse model. Proc Natl Acad Sci U S A. 2023 Aug 22;120(34):e2302676120.

35. Weiss SA, Fried I, Engel J, Jr, Sperling MR, Wong RKS, Nir Y, Staba RJ. Fast ripples reflect increased excitability that primes epileptiform spikes. Brain Commun. 2023;5(5).

36. Guth TA, Kunz L, Brandt A, Dümpelmann M, Klotz KA, Reinacher PC, Schulze-Bonhage A, Jacobs J, Schönberger J. Interictal spikes with and without high-frequency oscillation have different single-neuron correlates. Brain. 2021 Nov 29;144(10):3078–88.

